# In the chick embryo, estrogen can induce chromosomally male ZZ left gonad epithelial cells to form an ovarian cortex, which supports oogenesis

**DOI:** 10.1101/568691

**Authors:** Silvana Guioli, Debiao Zhao, Sunil Nandi, Michael Clinton, Robin Lovell-Badge

## Abstract

In chickens, as in most amniotes, the first morphogenetic event in ovary differentiation is the formation of two distinct domains: a steroidogenic core, the medulla, overlain by the germ cell niche, the cortex. This process normally starts immediately after sex determination in the ZW embryos, substantially before the progression of germ cells into meiosis. In order to address the extent to which the cortical domain depends on intrinsic or extrinsic factors, we generated models of gonadal intersex by mixing ZW (female) and ZZ (male) cells in gonadal chimeras, or by altering estrogen levels of ZZ and ZW embryos *in ovo*. We found that both chomosomally female and male cells can be induced to form the cortical domain and that this can form relatively independently of the phenotypic sex of the medulla as long as estrogen is provided. We also show that the cortex promoting activity of estrogen signalling is mediated via Estrogen Receptor alpha within the left gonad epithelium. Therefore, either a ZW or ZZ cortical domain can provide an adequate niche to allow the germ cells to progress into meiosis. However, the presence of a medulla with an “intersex” or male phenotype may compromise this progression, causing cortical germ cells to remain in an immature state in the embryo.

## Introduction

The gonads provide a paradigm in which to investigate organogenesis given that there are two possible outcomes, ovaries and testes, which arise from a common anlagen and cell lineages.

In the chick, gonadal sex specific morphological differences become grossly apparent from around day (D) 8 (HH34) (Hamburger and Hamilton, 1951) with the re-localisation of the germ cells to the appropriate domain. In chromosomally male (ZZ) embryos the germ cells become embedded within somatic cells to form the sex cords of the gonadal core or medulla, while in chromosomally female (ZW) embryos the germ cells aggregate at the periphery of the gonad and intermingle with somatic cells to form the cortex (Carlon and Stahl, 1985). In the embryonic phase of ovary differentiation, the germ cells organise in nests. It is only after hatching, when the cortex undergoes a major restructuring, in a process known as folliculogenesis, that the nests are broken and each germ cell becomes enveloped by layer(s) of granulosa cells. The embryonic cortex is therefore transient, but it is a common feature of most amniotes, including many mammals. It precedes the entry of germ cells into meiosis. Mouse and rat are exceptions to this rule, because their germ cells remain distributed in the whole gonad long after meiotic entry (Byskov, 1986; DeFalco and Capel, 2009).

The formation of the embryonic cortex is side dependent in the chick. This is due to the interaction of the canonical left-right asymmetry pathway with the gonadal development pathway via the transcription factor *Pitx2*, expressed asymmetrically in the left gonadal epithelium (Guioli and Lovell-Badge, 2007; Ishimaru et al., 2008; Rodriguez-Leon et al., 2008). During sex specific differentiation of the gonads, this asymmetry is ignored or overwritten in males, while it is exacerbated in females. Indeed, at sex determination, the medulla of both left and right embryonic ZW gonads differentiate into steroidogenic domains, but only the left gonad develops a cortex and proceeds to differentiate further into a functional ovary, while the right ovary remains just as a steroidogenic organ and later atrophies. Germ cells scattered in the medulla, in both the left and right ovaries, do not properly enter meiosis and are lost post hatching (Guioli et al., 2014). This asymmetry points to the gonadal epithelium as pivotal in the formation of the ovarian cortex and establishes the importance of the embryonic cortex for survival and maturation of the germ cells.

Recent chimera studies have demonstrated that somatic cells of the chick gonad medulla differentiate into either testicular cells or ovarian steroidogenic cells in a cell autonomous manner, i.e. dependent on their sex chromosome constitution (Zhao et al., 2010), however the importance of intrinsic sex chromosome identity in the formation of the cortical germ cell niche has yet to be addressed.

Like lower vertebrates and many mammals, although not the mouse, the embryonic ovaries of birds produce estrogen from the start of female specific differentiation. For chicken and some other vertebrate species, including some mammals, it has been shown that ectopic manipulation of estrogen levels at the time of sex determination can override the effects of chromosomal sex. Notably, blocking the activity of P450 aromatase, and as a consequence preventing the synthesis of estrogen, in ZW embryos before sex differentiation, results in female-to-male sex reversal. In these birds, both the left and right medulla start to express male markers, such as *SOX9*, in differentiating cords and the left epithelium fails to make a cortex. Conversely, injecting estrogen before sex differentiation in ZZ embryos results in male-to-female sex reversal. Although this phenotype is reported to revert to normal in the adult, in the embryo both medullas are reported to express *FOXL2* and *CYP19A1* and a cortex develops on the left side (Akazome and Mori, 1999; Bruggeman et al., 2002; Vaillant et al., 2003; Yang et al., 2008).

During chick sex determination, *Estrogen Receptor alpha* (*ERα*) is expressed in both the left and right medulla, but asymmetrically in the epithelium of the left gonad, (Andrews et al., 1997; Guioli and Lovell-Badge, 2007). This makes it a good candidate for the estrogen transducer, with the hypothesis that estrogen affects the differentiation of both medulla and cortex by acting on different cell types and different pathways. Furthermore, it suggests once again the pivotal role of the epithelium in the formation of the cortex.

In order to understand the process of embryonic cortex morphogenesis we investigated the importance of estrogen signalling in cortex differentiation in relation to the chromosomal sex of gonadal cells. By following the fate of mixed sex gonadal chimeras and of gonads derived from embryos with manipulated estrogen levels, we show that estrogen is the only signal necessary for cortex formation on the left side and that the cortex can be made of either ZW or ZZ cells without a bias. However, an intersex phenotype may compromise the progression of the germ cells to meiosis. Finally, we show that downregulating epithelial ERα is sufficient to severely affect cortex differentiation, indicating epithelial ERα as the relevant signal transducer.

## Results

### Modifying estrogen levels after the point of sex determination affects cortex formation without affecting the sex identity of the medulla

In order to understand the role of estrogen in cortex differentiation and the relationship between sex specific differentiation of cortex and medulla we altered estrogen levels beyond the time when sex reversal can be achieved (Bruggeman et al., 2002). To block/reduce estrogen levels we treated D7-7.5 (HH31) ZW embryos with the aromatase inhibitor fadrozole and repeated the administration every two days (ZW-Fa embryos) (Fig. 1). Gonads recovered at D10 (HH36) showed a female medulla as expected, with no sign of masculinisation, as no male markers such as SOX9 were identified by immunostaining, similar to the ZW wildtype. However, the cortical domain of the left ovaries appeared reduced in thickness compared to controls and contained fewer germ cells (Fig. 1 C). ZW left ovaries collected at D17 (HH43) were morphologically much smaller compared to ZW controls (suppl. Fig. 1), but still had a cortical domain. However, this was generally limited to the central part of the ovary (Fig. 1 G).

**Fig. 1.**
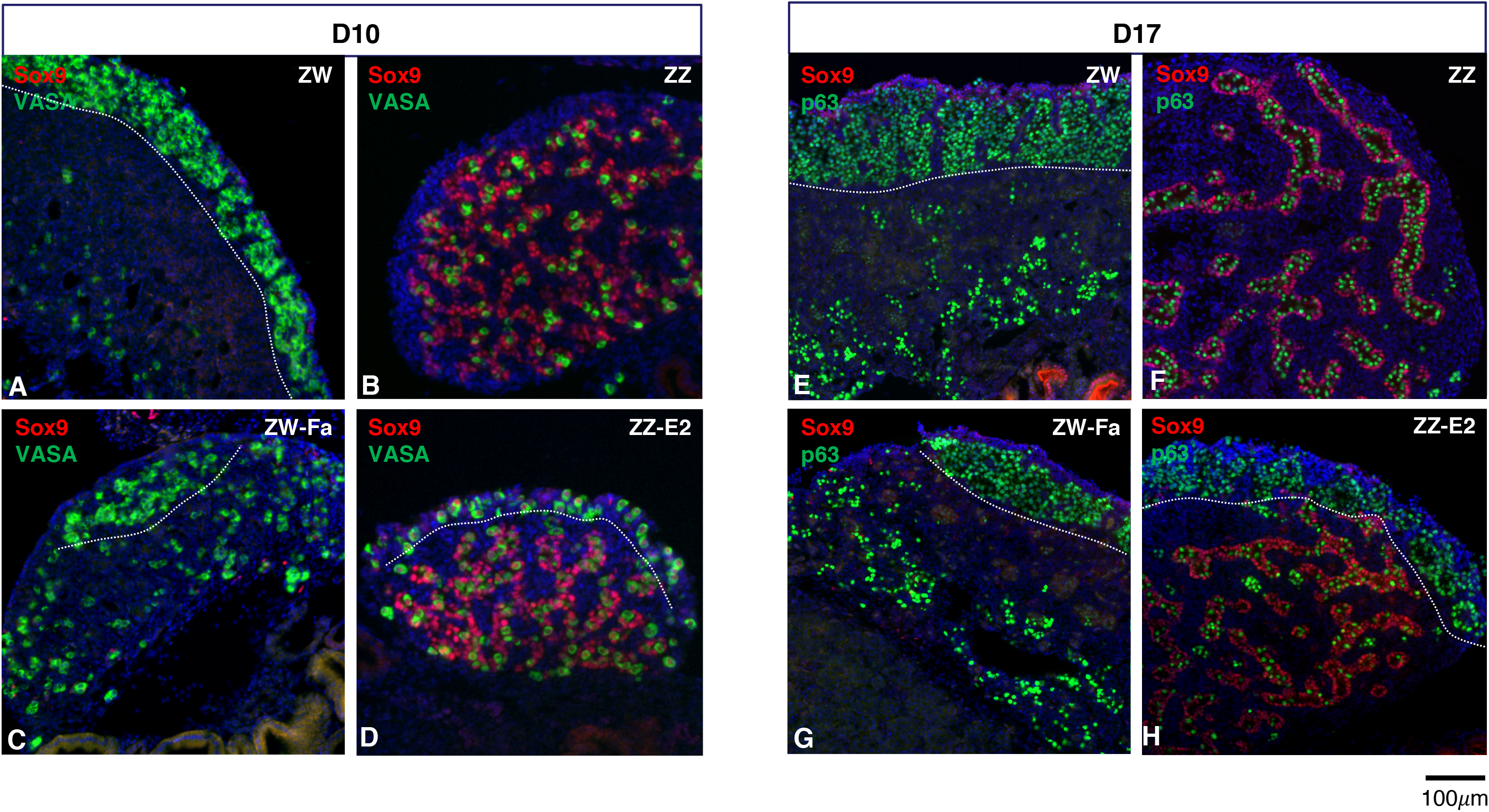
Perturbing estrogen levels at embryonic D7-7.5 (HH31) affects cortex formation in ZW and ZZ embryos. Sections from left gonads at D10 (A-D) or D17 (E-H) double-stained for the Sertoli marker SOX9 (red) and a germ cell marker: VASA or P63 (green). (A, E) ZW controls; (B, F) ZZ controls; (C, G) ZW gonads treated with Fadrozole (ZW-Fa); (D, H) ZZ gonads treated with ß-estradiol (ZZ-E2). Decreasing estrogen in ZW embryos after sex determination compromises the differentiation of the ovarian cortex; adding ß-estradiol in ZZ embryos after sex determination induces the formation of a cortex on top of a male medulla. White dotted lines highlight the cortex-medulla border.

To upregulate estrogen in ZZ embryos after sex determination, we injected β-estradiol *in ovo* at D7-7.5 (HH31) (ZZ-E2 embryos) (Fig. 1). The resulting ZZ left gonads collected at D10 (HH36) comprised a male medulla containing cords made of SOX9-positive somatic cells and germ cells, overlain by a narrow cortex-like domain (Fig. 1 D). Left and right gonads recovered at D17 (HH43) showed a more striking phenotype, with a well-developed male medulla on both sides and a quite thick cortical domain on the left side, containing germ cell nests (Fig. 1 H). Similar results were obtained when β-estradiol was injected much later (i.e. at D9 (HH35), suppl. Fig. 2).

**Fig. 2.**
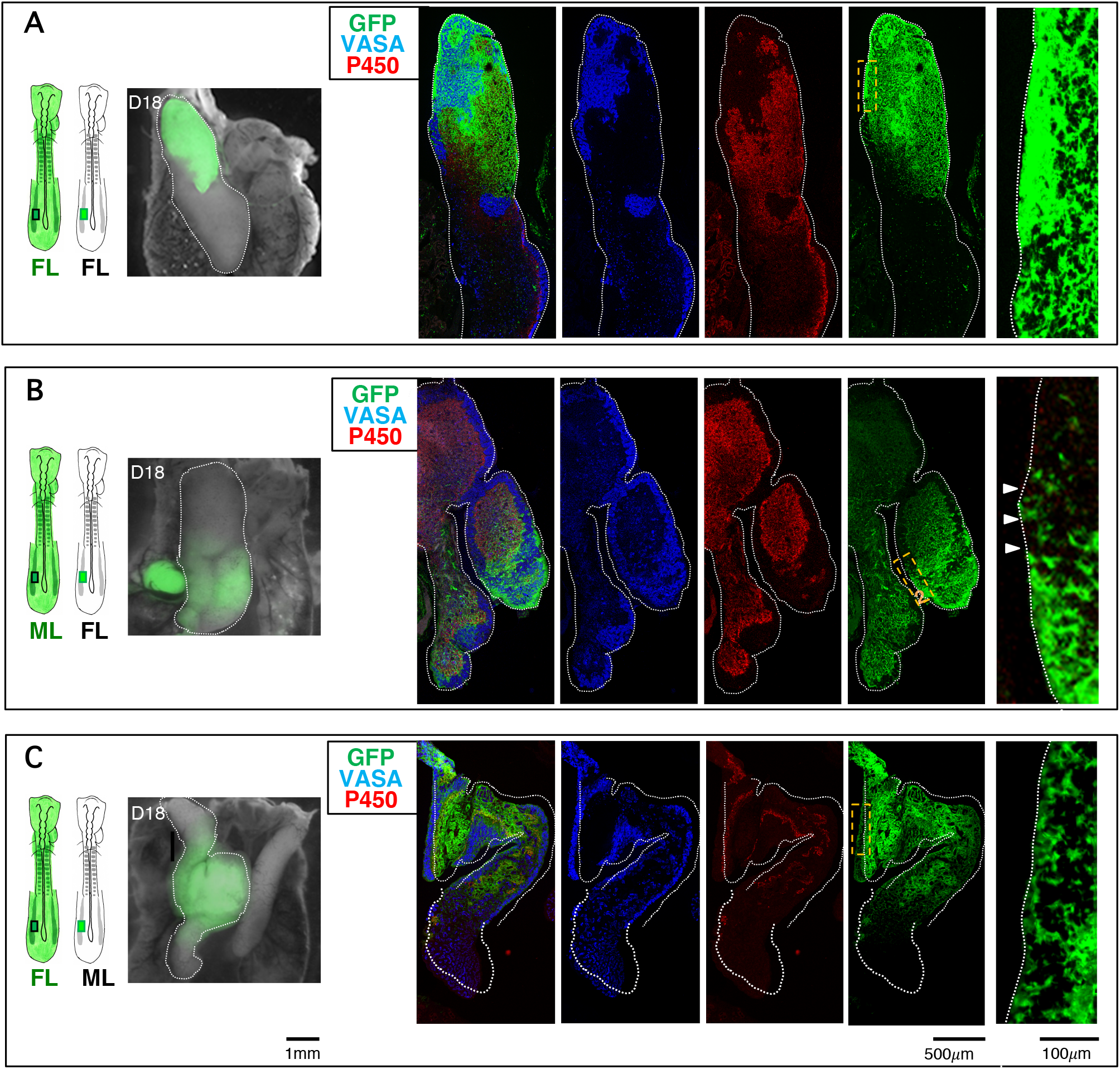
ZW and ZZ cells can both contribute to the somatic component of the cortex domain. D18 gonadal chimeras generated by transplanting posterior lateral plate/intermediate mesoderm from donor GFP embryos (green) into recipient wildtype embryos at D1.5-2 (HH10). In each panel (A-C) from left to right: schematic of the transplant; whole-mount image of the resulting gonads dissected at D18 (HH44); immunofluorescent images of sections from the left chimeric gonads showing the cells positive for the germ cell marker VASA (blue), P450aromatase (red) and GFP (green); on the far right side, high magnification of the GFP pattern within the cortical area in the orange dashed box. (A) Female-Left (FL) to female-Left (FL) chimeric ovary. (B) Male-Left (ML) to Female-Left (FL) chimeric ovary. (C) Female-Left (FL) to Male-Left (ML) chimeric ovotestis. The ZZ cells can contribute the somatic component of the cortex without sex bias. White dotted lines highlight the chimeric left gonad borders. Arrow heads show areas of intermingling of ZW and ZZ epithelial cells.

These results show that the early differentiation of the cortex can be independent of the sex of the medulla and points to estrogen as the key inducer of this process.

### ZW and ZZ cells can contribute to the cortical domain in mixed sex gonadal chimeras

It has previously been shown that, in the chick, the medulla of the gonad differentiates in a cell autonomous manner, depending on the chromosomal sex of the component cells (Zhao et al., 2010). As the differentiating embryonic ovary is composed of two distinct sex specific functional domains: a steroidogenic medulla and a cortical domain essential for germ cells maturation, it remains to be assessed how important chromosomal sex identity is for the formation and differentiation of a proper ovarian germ cell niche. The estrogen manipulation experiments have demonstrated that a cortex can form in a ZZ embryo, however it is not clear that this is possible under normal physiological conditions. To address this issue, we generated gonadal chimeras comprised of male and female cells by transplanting D2 (HH10) lateral plate/intermediate mesoderm from the left side of donor GFP embryos into matching age and same side recipient embryos of the same or opposite sex, *in ovo*. The manipulated embryos were re-incubated and the gonads were collected at D18 (HH44) (Fig. 2).

The resulting manipulated embryos contain a host derived right gonad and a left gonad containing somatic cells of host and donor origin. The germ cells, however, are always host derived on both sides, as they segregate to the germinal crescent of the embryo at stage HH5 and later migrate to the gonad via the bloodstream (Ginsburg and Eyal-Giladi, 1987; Karagenc et al., 1996; Nakamura et al., 1988).

The chimeric gonads derived from left-to-left transplants between ZW embryos (FL-FL, same-sex chimeras) were ovaries composed of a medulla and a cortex similar to those in a normal left ovary. In the donor enriched area, the cortical epithelial and sub-epithelial somatic cells were mainly contributed by GFP donor cells (Fig. 2 A).

Fig. 2 B shows a typical chimeric left ovary from a ZW embryo containing donor cells derived from ZZ left mesoderm (ML-FL mixed-sex chimera). This ovary displays a cortex along the entire length. The area enriched in donor tissue was composed of a medulla containing host (ZW), P450arom positive steroidogenic cells, surrounded by host (ZW, GFP negative) and donor (ZZ, GFP positive) derived interstitial cells, overlain by a cortex formed by GFP donor (ZZ) cells. These areas were continuous with areas poor in donor cells, whose medulla and cortex were mainly host derived. At the junction between the two areas, intermingling of host and donor cells of the cortex was evident. Transplants of ZW donor tissue into ZZ hosts (FL-ML mixed-sex chimeras) often produced chimeric ovotestes. Fig. 2 C shows one such example. The ovarian portion of the chimera comprised a female medulla containing P450arom positive steroidogenic cells derived from the donor (ZW, GFP positive cells) as expected, overlain by a cortex made of both host (ZZ) and donor (ZW) somatic cells.

We concluded that both ZZ and ZW cells can normally contribute the somatic component of the embryonic cortex without a sex bias and therefore independently of their chromosomal sex.

### Meiotic entry of cortical germ cells is severely compromised in gonads with a masculinised medulla

The mitotic to meiotic switch is quite asynchronous in the chicken ovarian cortex, with many germ cells entering meiosis by D12 (HH38) and the majority initiating the synaptic process by D16 (HH42) (de Melo Bernardo et al., 2015).

We followed germ cell meiotic progression in D17 (HH43) ovaries from ZW embryos subject to late fadrozole treatment (ZW-Fa) and from ZZ embryos subject to late estradiol treatment (ZZ-E2), by analysing the expression of the double strand breaks (DSB) marker γH2AX and the synapsis marker SYCP3 (Guioli et al., 2012; Smith et al., 2008). At this stage, both markers are normally expressed in most germ cells (Fig. 3). Similarly, many germ cells expressed both markers in the estradiol treated ZW left gonads. In fadrozole treated ZW left gonads the cortex contained germ cells positive for γH2AX and SYCP3, although visibly fewer than in the ZW control. Meiosis was most affected in estradiol treated ZZ left gonads, composed of a medulla organised in testicular cords, overlain by a cortical domain made of ZZ somatic cells and ZZ germ cells: some cortical germ cells expressed γH2AX but none were positive for SYCP3 as assessed by immunostaining (Fig. 3).

**Fig. 3.**
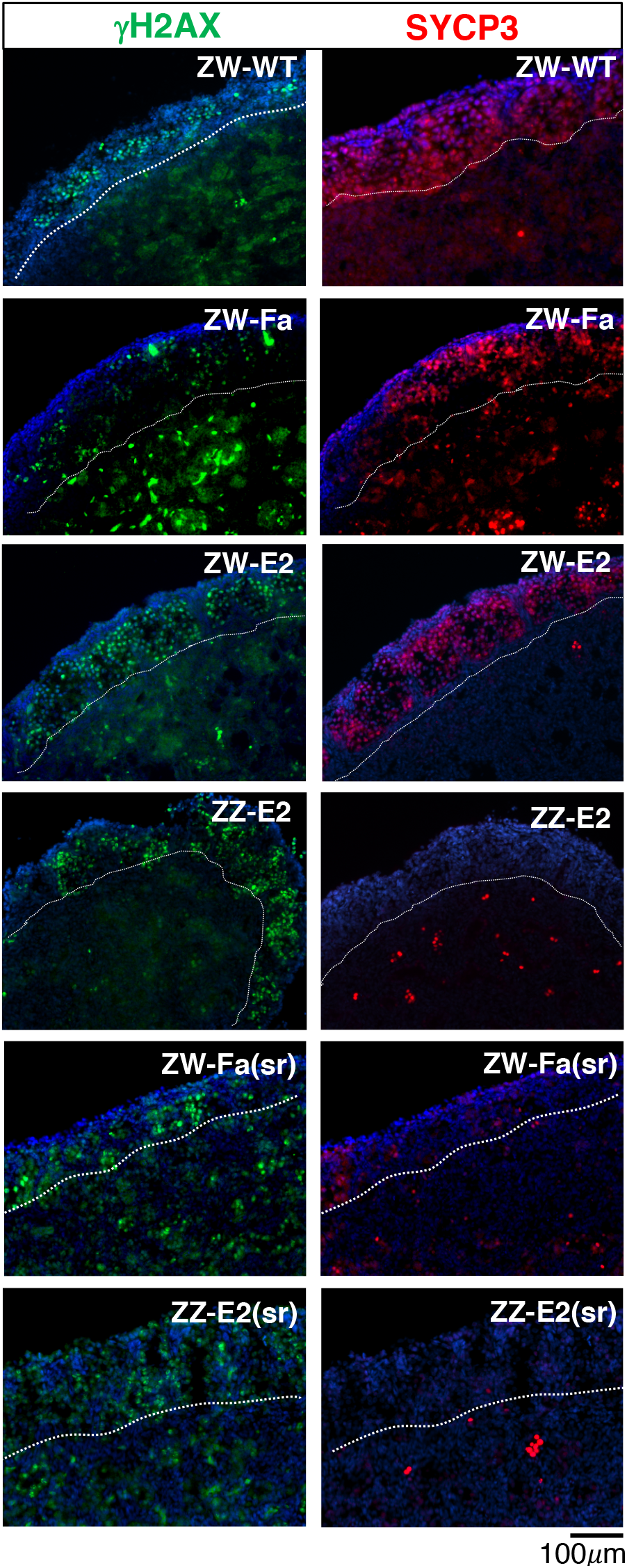
Induction of meiosis is compromised in the cortical germ cells of D17 embryos subject to estrogen level alterations. D17 (HH43) left gonad sections immunostained for γH2AX (green) or SYCP3 (red). ZW-WT: ZW wildtype gonad. ZW-Fa: ZW gonad treated with fadrozole from D7-7.5 (HH31); ZW-E2: ZW gonad treated with ß-estradiol at D7-7.5; ZZ-E2: ZZ gonad treated with ß-estradiol at D7-7.5; ZW-Fa(sr): ZW, partially sex reversed gonad, treated with Fadrozole at D4; ZZ-E2(sr): ZZ, partially sex reversed gonad, treated with ß-estradiol at D4. In ZW-Fa and ZW-E2 many germ cells express SYCP3, although SYCP3 appears to be less widespread in ZW-Fa than ZW control. In ZZ-E2(sr), ZZ-E2 and ZW-Fa(sr) some germ cells express γH2AX but none or very few express SYCP3.

To investigate the importance of germ cell chromosomal sex for the correct initiation of meiosis in the cortex, we generated a series of testis-to-ovary reversed ZZ embryos and a series of ovary-to-testis reversed ZW embryos, by *in ovo* injection of estradiol or fadrozole, respectively, at D4 (HH23) (ZZ-E2(sr) and ZW-Fa(sr) embryos). From both experiments, we obtained D17 embryos with incomplete gonadal sex reversal which typically showed an “ovotestis” on the left. The inner part of the medulla contained SOX9-positive cells, organised in cords like structures, while the outer areas contained more FOXL2 positive cells (data not shown). The medulla was generally overlain by a cortex like structure (Fig. 3). Importantly, in ZZ-E2(sr) embryos the primordial germ cells (PGCs) are male (ZZ) while in ZW-Fa(sr) embryos they are female (ZW). Synapsis was compromised in both these models, because in both cases some cortical germ cells expressed γH2AX but none or very few expressed SYCP3 by immunostaining (Fig. 3).

*STRA8* is a major factor involved in meiotic initiation and should be expressed in germ cells of the left cortex starting from D12 (HH38) (Smith et al., 2008). We analysed the *in situ* expression pattern of *STRA8* in the cortex of ZZ embryos exposed to late estradiol treatment. These were negative for *STRA8* at D14 (HH40) and showed only a few very faint patches at D17 (HH43), compared to the cortex of normal ZW controls and estradiol treated ZW embryos. In contrast, the D17 (HH43) cortex of ZW embryos exposed to late fadrozole treatment was positive, although at lower levels than ZW controls (Fig. 4 A).

**Fig. 4.**
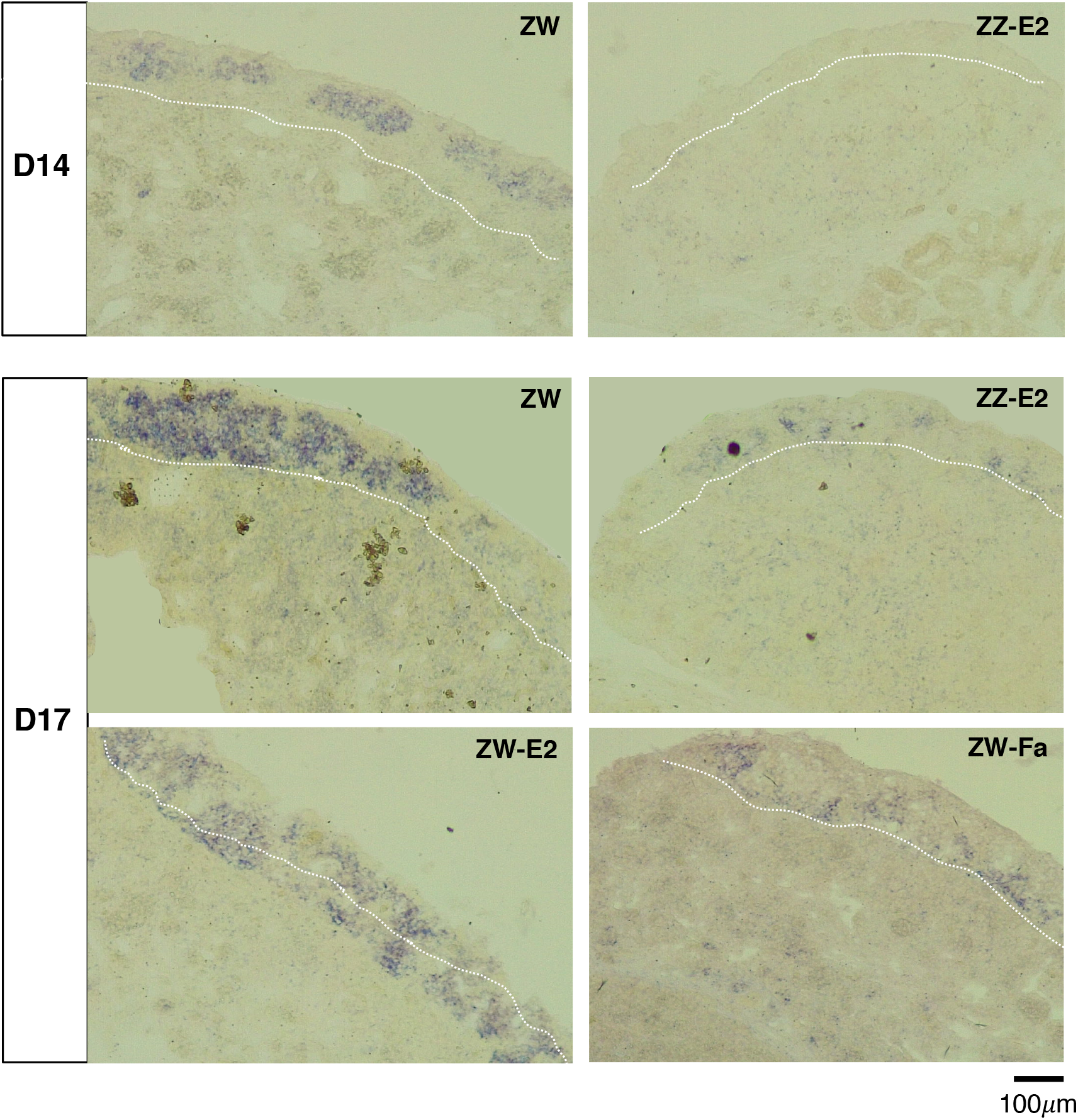
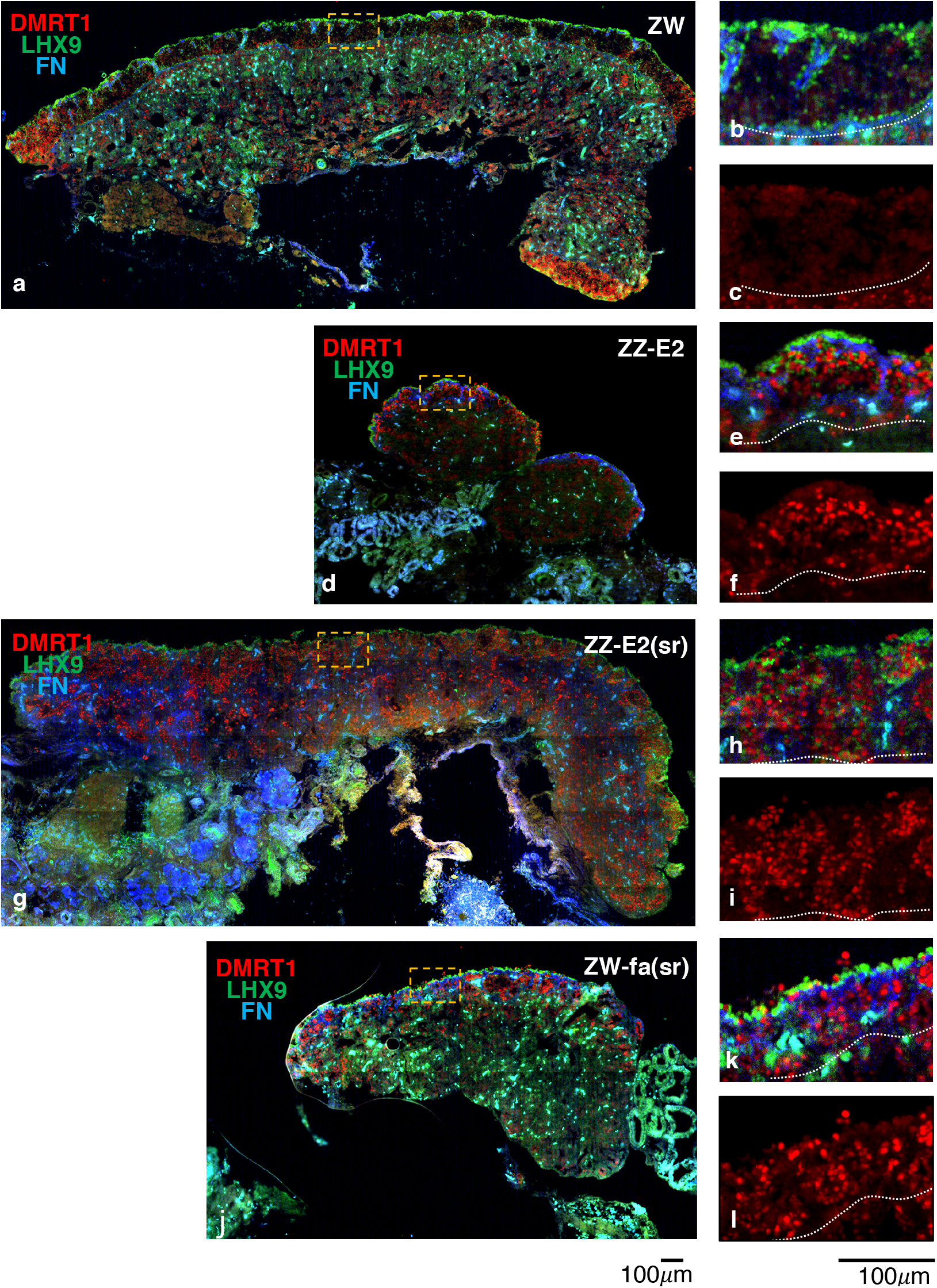
The expression pattern of mitotic-meiotic switch markers is affected in the cortical germ cells of embryos subject to estrogen levels alteration. (A) RNA *in situ* analysis of *STRA8* expression on sections from the left gonad of D14 (HH40) and D17 (HH43) embryos. ZW: female wildtype control; ZZ-E2: ZZ, ß-estradiol treated at D7-7.5 (HH31); ZW-E2: ZW, ß-estradiol treated at D7-7.5 (HH31); ZW-Fa: ZW, Fadrozole treated at D7-7.5 (HH31). *STRA8* expression is severely compromised in the gonadal cortical germ cells from ZZ embryos exposed to ß-estradiol. (B) Immunofluorescent detection of DMRT1 (red), LHX9 (green), and fibronectin (FN) (blue) on cryostat sections from the left gonad of D17 (HH43) embryos. LHX9 marks cortical somatic cells and and FN highlights the cortex-medulla border cells, respectively (Guioli and Lovell-Badge, 2007). DMRT1 marks the cortical germ cells, not the somatic cortical cells (LHX9 positive). (a-c) ZW: female wildtype control; orange dashed area in (a) enlarged in (b-c), LHX9+FN+DMRT1 and DMRT1 only, respectively. (d-f) ZZ-E2: ZZ, treated with ß-estradiol at D7-7.5 (HH31); orange dashed area in (d) enlarged in (e-f), LHX9+FN+DMRT1 and DMRT1 only, respectively. (g-i) ZZ-E2(sr): ZZ, treated with ß-estradiol at D4 (HH23) (partially sex reversed); orange dashed area in (g) enlarged in (h-i), LHX9+FN+DMRT1 and DMRT1 only, respectively; (j-l) ZW-Fa(sr): ZW, treated with fadrozole at D4 (HH23) (partially sex reversed); orange dashed area in (j) enlarged in (k-l), LHX9+FN+DMRT1 and DMRT1 only, respectively. In the control female DMRT1 is only present in germ cells at the gonad poles. In ZZ-E2, ZZ-E2(sr) and ZW-Fa(sr) DMRT1 is expressed in many germ cells across the cortex. White dotted lines show the cortex-medulla border.

We also checked the expression of *DMRT1*, a known key factor involved in the mitotic-meiotic switch in mouse (Matson et al., 2010; Zarkower, 2013). In chick, as in the mouse, *DMRT1* is normally expressed in the proliferating germ cells in both differentiating ovary and testis and it is downregulated soon after meiosis starts (Guioli et al., 2014; Omotehara et al., 2014). As shown in Fig. 4B, DMRT1 is downregulated by D17 (HH43) in most germ cells of a wildtype ovary. DMRT1-positive cells were only observed at the poles of the control ovary where the maturation of germ cells is delayed (de Melo Bernardo et al., 2015). In the experimental samples ZZ-E2, ZZ-E2(sr) and ZW-Fa(sr) that were severely compromised for SYCP3 expression, numerous germ cells throughout the cortex were strongly positive for DMRT1 (Fig. 4 B).

### A cortical domain contributed by ZZ male somatic cells provides a proper niche for ZW and ZZ germ cell meiotic entry

In order to assess if a cortical domain made of ZZ somatic cells can sustain the progression of the germ cells into meiosis in physiological conditions, we analysed the expression of the meiotic markers in D18 (HH44) mixed-sex gonadal chimeras (Fig. 5). In the same-sex female left chimeric ovary (FL-FL) most germ cells localised along the entire cortex were positive for both γH2AX and SYCP3 as expected (Fig. 5 A-B). In ML-FL mixed-sex left chimeric ovary, where the germ cells are female (ZW), the pattern of cortical PGCs was similar to the FL-FL controls. This means that SYCP3 was expressed in most PGCs throughout the entire cortex, both in areas mainly contributed by host female (ZW) somatic cells and in areas mostly contributed by donor male (ZZ) somatic cells, demonstrating that the latter are competent to form a proper cortical germ cell (oocyte) niche (Fig. 5 C-D). However, in FL-ML left chimeric ovotestis, where the germ cells are male (ZZ), the results were more complex. Within the ovarian portion the cortical germ cells showed a variable pattern depending on localisation. In some cortical areas, many germ cells were found positive for γH2AX and SYCP3, while in other areas few or no germ cells were positive for these markers (Fig. 5 E-G). The fact that their ability to enter meiosis appears to be site dependent, indicates that both female and male germ cell chromosomal sex identity is compatible with starting the female meiotic program in the embryo.

**Fig. 5.**
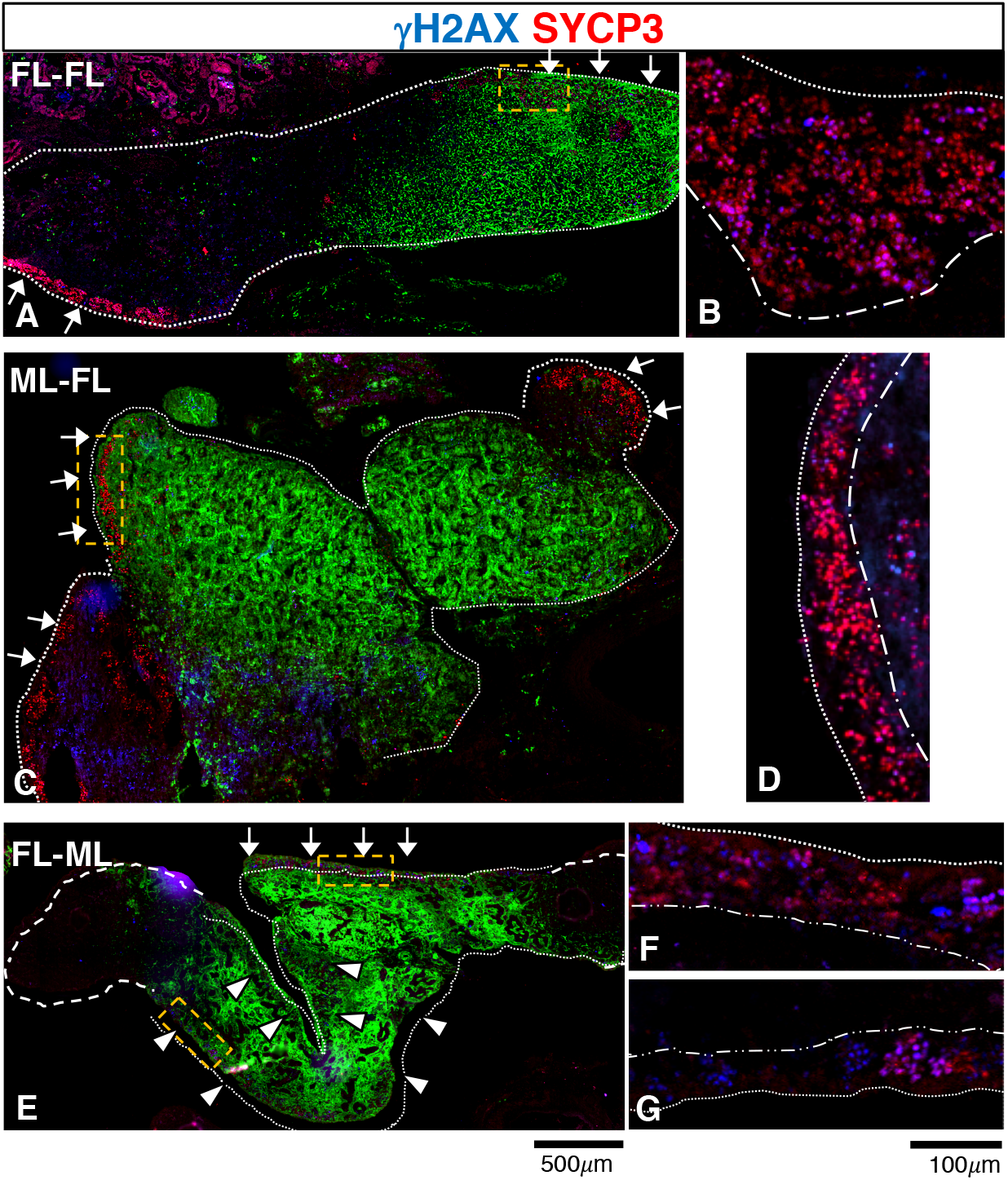
In mixed sex gonadal chimera ZZ somatic cells provide an adequate niche for progression of cortical germ cell into meiosis. Sections from chimeric left gonads at D18 (HH44) immunostained for γH2AX (blue) and SYCP3 (red). Donor cells are GFP positive (green). (A) female left into female left (FL-FL) control chimeric ovary; (C) male left into female left (ML-FL) mixed-sex chimeric ovary (E) female left into male left (FL-ML) mixed sex chimeric ovotestis; (B), (D) and (F-G) are high magnification of orange dashed areas in (A), (C) and (E) respectively. In FL-FL and in ML-FL chimeric ovaries most cortical germ cells express SYCP3. In the FL-ML chimeric ovotestis, SYCP3 is expressed in some cortical germ cells depending on localisation. Arrows indicate the cortical areas where many SYCP3 cells are present, arrow heads indicate the areas with very few SYCP3 cells. White dotted lines highlight the borders of the chimeric ovaries or ovarian domain of ovotestis, white dashed lines highlight the testis portion of the ovotestis. White dashed-dotted lines in enlargement panels show the cortex-medulla border. In (B), (D), (F), (G) enlargement panels GFP has been omitted for better visualisation of the meiotic markers pattern.

### ERα downregulation in the ZW left gonad epithelium disrupts cortex differentiation

At sex determination, *ERα* is asymmetrically expressed in both ZW and ZZ embryos, being restricted to the epithelium of the left gonad although present in both medullas (Andrews et al., 1997; Guioli et al., 2014). This pattern makes it a candidate estrogen signal transducer in promoting cortical differentiation. In this study, we showed that the ZZ left gonad is sensitive to estrogen beyond the initiation of male specific differentiation, implying that an estrogen transducer should be expressed asymmetrically in the differentiating testis. However, this had not been specifically addressed in previous studies. To address this issue we performed a timecourse analysis of ERα during testis differentiation by immunostaining. Its expression was clearly detected in the epithelium of the left ZZ gonad at D7 (HH31). Although weaker than in the differentiating ovary, the staining was maintained in the epithelium of the left testis until at least D12 (Suppl. Fig. 3). This indicates that the left epithelium of the ZZ developing testis retains the potential to quickly respond to estrogen, even though it would not normally be exposed to it at any significant level.

In order to knock down the activity of ERα in the epithelial cells of the left ovary, a dominant negative form of ERα (cERα524),cloned into a Tet-ON plasmid for conditional gene expression, was transfected into the left gonadal epithelium at D2.5 (HH15-17) by *in ovo* electroporation of dorsal coelomic epithelium on the left side (Guioli et al., 2007). The DNA was co-electroporated with other plasmids of the Tet-ON system, including T2TP for expression of the Tol2 transposase and pT2K-CAG-rTA-M2, the doxycylin-inducible activator (Watanabe et al., 2007) (Fig. 7A and M&M). cERα524 expression was induced at D4.5 (HH25) or D5.5 (HH27) and the screening was carried out at D10 (HH36) based on the EGFP reporter expression, as a measure of the quality of transfection. Electroporation typically results in mosaic transfection which varies from sample to sample. ZW embryos displaying high levels of EGFP (e.g. Fig. 6 B-C) were processed for immunostaining with markers specific to different cell types. A range of phenotypes were identified within the cortex, ranging from thinning, to lack of a proper cortex along most of the ovary. The most severe phenotype was observed in ovaries exposed to doxycycline at the earliest time point. An example is shown in Fig. 6 B, E, H, M. In this ovary, most of the medulla highlighted by P450 aromatase was overlain by a simple epithelium, with the exception of the central part, which displayed some stratification and contained a small aggregate of germ cells (Fig. 6 E, H). An example of a weaker phenotype is shown in Fig. 6 C, F, I, N. The cortex was quite irregular in size and markedly thinner in areas rich in EGFP. Germ cells density along the cortex varied accordingly to the cortex size in the electroporated samples (Fig. 6 I). Although by D10 (HH36) many EGFP epithelial derived cells were found in the medulla, the medulla maintained an ovarian identity, being positive for P450 (Fig. 6 E-F) and negative for SOX9 (data not shown). No other obvious phenotype was observed in the medulla. All samples still expressed *Pitx2* within the left epithelium as seen in the normal controls (Fig. 6 L-N).

**Fig. 6.**
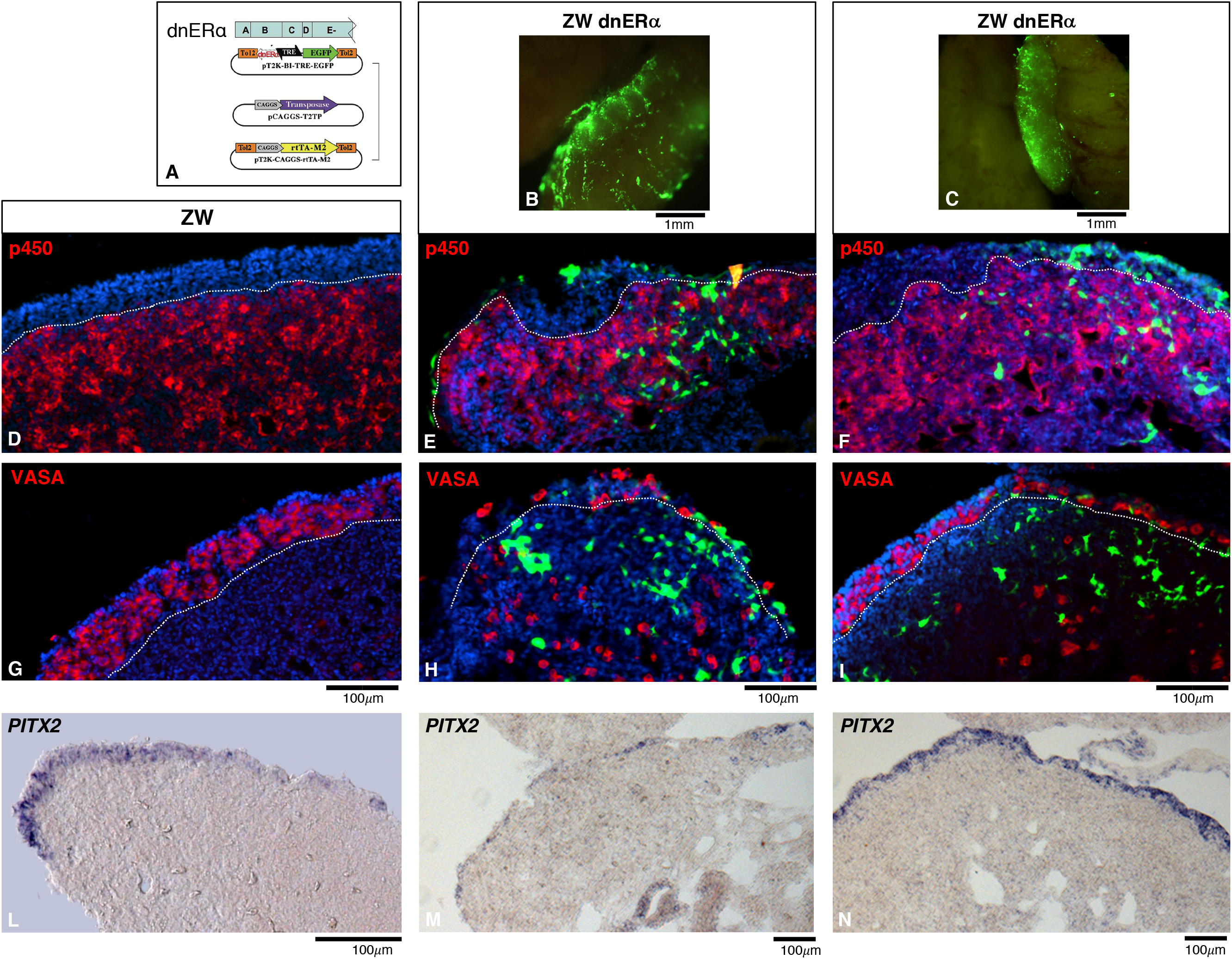
Suppression of epithelial ERα activity disrupts cortex differentiation. (A) Schematic of the inducible TET-ON plasmid system used to express a dominant/negative isoform of ERα (dnERα) in the left gonad epithelium. (B-C) Whole mount images of two ZW left gonads electroporated at D2.5 (HH15-17) and screened at D10 (HH36). Targeted cells are highlighted by expression of an EGFP reporter (green). (D-I) Fluorescence images of D10 (HH36) gonad sections immunostained for P450aromatase (P450) (red) or the germ cell marker VASA (red); (D, G) ZW control; (E, H) and (F, I) ZW electroporated gonads shown in (B) and (C), respectively. The cortex is severely compromised by downregulation of ERα activity. (L-N) *PITX2* RNA in situ expression pattern: (L) ZW control, (M) and (N) ZW electroporated gonads shown in (B) and (C), respectively. *PITX2* expression is not affected by downregulation of ERα in the left epithelium. White dotted lines show the cortex/medulla border.

To make sure that the data observed were due to specific knock down of ERα activity we performed an alternative set of experiments aiming to suppress ERα via RNAi. Supplementary Fig. 4 shows a left ovary screened at D10 (HH36) and analysed for germ cell localisation and cortical structure. As with the phenotype obtained with cERα524, the ER*α* shRNA targeted left gonad had a normal female medulla similar to the ZW wildtype control, but a severely compromised cortex in correspondence to areas enriched in EGFP cells.

## Discussion

In recent years great advances have been made uncovering key sex regulators that control the commitment to the male and female programmes, mainly based on work carried out in mammals, primarily in the mouse.

However ovarian differentiation is still poorly understood. One of the main reason is that mouse ovarian morphogenetic changes, including the formation of a cortex, are much delayed to the perinatal period, making the discovery of female sex promoting factors more challenging. Moreover, the insensitivity to estrogen in the mouse embryo may reflect the evolution of a different hierarchy of early regulators, compared even to other mammals. One example of this is *Foxl2* which is critical for ovary differentiation early in gonadal development in the goat, but not until after birth in the mouse (Boulanger et al., 2014; Schmidt et al., 2004; Uda et al., 2004; Uhlenhaut et al., 2009).

The chicken ZW gonad produces estrogen and differentiates into an ovary with a clear cortex and medulla soon after the female molecular pathway initiates, making it a good model system to address early ovarian differentiation. In this study, we focused on cortical morphogenesis with the aim of understanding the differentiation of the female (i.e. oogenic) germ cell niche.

Estrogen signalling is an important factor in chicken female sex determination, as it can override the influence of genetic sex if perturbed at the time of sex determination (Yang et al., 2008). This holds true in lower vertebrates and some mammals including marsupials (Coveney et al., 2001; Guioli et al., 2014). However, its potential role as anti-testis and ovarian promoting factor are still ambiguous. To explore its action in promoting cortex formation and to understand if this process can be dissociated from female specification of the medulla, we observed the effects of manipulating estrogen levels after D7 (HH31), when sex reversal is no longer achievable and medulla differentiation is consistent with genetic sex. We found that estrogen injection in ZZ embryos generates an “intersex” gonad composed of a male medulla overlain by a female cortex, while downregulation of estrogen in ZW embryos perturbed proper cortex formation, maintaining a female medulla. In parallel we generated a series of male:female gonadal chimeras and observed that even the presence of a small aromatase positive medullary domain was sufficient to induce cortex formation in the left gonad.

These results show that the cortex domain can, at least initially, differentiate independently from the sex of the medulla if estrogen is provided, even in small quantities. This makes estrogen the major promoting signal for cortex development and the left epithelium a naïve tissue capable of responding to any sex environment hormonal influence.

Therefore, early cortical formation depends mainly on the presence of promoting signals, notably estrogen, normally produced in the ovarian medulla, rather than on the lack of any antagonistic signals, produced, for example, in the male medulla.

Moreover, the analysis of same-sex and mixed-sex left gonadal chimeras showed that under physiological conditions, ZZ and ZW cells can become part of a cortex without a sex bias and that a cortex composed of ZZ somatic cells can provide a proper oogenic niche like the female counterpart. In both cases, after an initial phase of proliferation, the germ cells entered meiosis at the appropriate time, excluding the involvement of W or Z specific genes in the control of early cortical development. It is known that *Dmrt1*, a current candidate as a male primary sex determinant in chicken (Guioli et al., 2014; Lambeth et al., 2014; Smith et al., 2009), is expressed symmetrically in the medulla, but also asymmetrically in the gonad epithelium of ZZ and ZW embryos at the time of sex determination (Guioli and Lovell-Badge, 2007; Omotehara et al., 2014). As *Dmrt1* is not subject to dosage compensation (McQueen and Clinton, 2009), our chimeras show that any differences in *Dmrt1* expression levels due to gene dosage between female and male somatic cells does not impact cortex formation. Therefore, although the cortex is a distinct female specific domain of the ovary with somatic cells providing a niche for oogenesis, the chromosomal sex of these cortical somatic cells is irrelevant. This is in contrast to the somatic cells of the testis cords, which have to be ZZ in order to provide a functional niche for spermatogenesis. Thus, one cell type, the somatic cells of the medulla which differentiate in a cell autonomous manner strictly linked to their chromosomal sex (Zhao et al., 2010), drive the sex specific differentiation of the entire organ.

However, we did observe that in the left gonad of ZZ embryos subject to late estradiol treatment and in ovotestes commonly obtained from female to male (FL-ML) chimeras, cortical germ cells progression into meiosis was compromised, as SYCP3, a marker of the synaptic process was absent or expressed in very few cells at D17-18 of embryogenesis. Although in these severely compromised models the PGCs were genotypically male, there is evidence that this failure is not a result of the germ cell genotype. For example, although meiosis was compromised in FL-ML chimeras, localised areas with SYCP3-positive ZZ germ cells were found, suggesting that the ZZ germ cells are capable of initiating meiosis in the cortex, but they do so on the basis of their localisation within the cortical domain. Moreover, left ovotestes, generated by manipulation of estrogen levels in either D4 ZZ or ZW embryos, both contained cortical germ cells with compromised SYCP3 expression. This therefore occurs independently of the germ cell chromosomal sex.

In chicken, as in the mouse, *STRA8* is a key factor for germ cell entry into meiosis (Bowles et al., 2016; Koubova et al., 2014; Smith et al., 2008; Yu et al., 2013). In our SYCP3 compromised models *STRA8* was almost undetectable throughout development, while DMRT1, which should be downregulated at meiosis, was still present in many D17 (HH43) germ cells. This shows that at D17 (HH43) these cortical domains contained germ cells that have not progressed properly into meiosis, suggesting meiotic arrest or at least a severe delay.

The mitotic-meiotic switch is a very complex process, including many germ cell intrinsic and extrinsic factors which may act as promoters or suppressors of meiosis that are not yet completely elucidated. Our results suggest that extrinsic factors outside the cortex must be involved. It has been shown that FGF9 secreted by the Sertoli cells antagonises meiotic entry via activation of Nodal-Smad signaling in mouse spermatogonia (Tassinari et al., 2015) and in chicken ovary bFGF was found to act as a suppressor of meiosis and as a mitogenic factor (He et al., 2012). The proximity of a male medulla in the ovotestes may antagonise meiotic entry via FGF signalling or other male driven signalling. Alternatively, meiosis may be impaired by the lack or deficiency of some female inducer signals from the medulla, or both.

In humans, disorders of sexual development (DSD) are associated with an increased risk of type II germ cell cancer which are derived from two types of *in situ* neoplastic lesions: carcinoma *in situ* (CIS) or gonadoblastoma, depending on the supporting cells being Sertoli cells or granulosa cells, respectively (Hersmus et al., 2017). The observation of mitotic and meiotic signals within CIS cells, together with the inability of these cells to ever enter meiosis, points to a dysfunctional mitotic:meiotic switch that could provide the genetic instability initiating the tumour (Cools et al., 2006; Jorgensen et al., 2013). The long-standing hypothesis is that this process is initiated in fetal life, where a gonad with an intersex phenotype provides a sexually confused niche which can result in the delay or arrest of germ cell development. If these cells are not eliminated they may retain the embryonic phenotype and later transform (Jorgensen et al., 2013).

These human studies support the idea that the meiotic phenotype in our intersex chick models is due to the sexually confused environment and predict that if these germ cells are not later eliminated, they could become transformed into tumour cells. Therefore, the estrogen manipulated chick models might even provide an important system in which to study the origin and dynamics of neoplastic transformation of germ cells.

We have previously proposed that estrogen signalling could be transduced via ERα on the basis of the RNA expression pattern at sex determination (Andrews et al., 1997; Guioli and Lovell-Badge, 2007). The symmetric expression in left and right medulla and the asymmetric expression in the gonad left epithelium of both female and male embryos suggested that estrogen, via ERα, may have multiple roles in promoting ovary differentiation which affects both the differentiation of the medulla and the cortex. However the protein pattern in the male has always remained elusive, making it difficult to explain how the radiolabelled estradiol injected in the embryo found to be binding both female and male gonad at D5-10 may do so via ERα (Gasc, 1980). In the present study we were able to show that some levels of ERα protein are present in the male gonad at sex determination in a pattern which reflects the RNA expression, indicating that both ZW and ZZ gonad could quickly respond via ERα to estrogen stimulation. Moreover, the maintenance of expression within the left epithelium in the differentiating testis shows that the epithelium remains exquisitely sensitive to estrogen and able to rapidly respond to the hormone even several days into the male differentiation pathway.

In agreement with predictions, the knock down of ERα activity in the epithelial cells of the left ZW gonad has resulted in the impairment of cortical development, the most severe cases displaying areas devoid of a cortex and overlain by a simple epithelium, similar to the phenotype observed in the D10 embryos treated with fadrozole from D7-7.5 (HH31). The downregulation only affected the cortical domain without affecting the sex identity of the medulla. Moreover, the cortical phenotype was not linked to variation in *PITX2* expression within the left epithelium. All together these data indicate ERα as the main transducer of estrogen signalling in cortex formation and that this process strictly depends on its activity within the left epithelial cells of the gonad.

ER is a complex molecule. It has been shown in different systems that upon estrogen activation, it can have non-genomic and genomic activities. It can rapidly respond at the plasma membrane by triggering signalling pathways and it can mediate transcriptional regulation within the nucleus (Hamilton et al., 2017). Our future work will be addressing the mechanism of action of ERα in promoting cortical development upon estrogen stimulation, through the identification of the downstream targets of its action via transcriptomics approaches.

In summary, our data suggest a model whereby the gonads, at the time of somatic sex specific differentiation, are formed of two distinct domains: the epithelium and the medulla. Left and right medulla differentiates along the male or female pathway depending on whether the cells comprising it are ZW or ZZ. On the other hand, the somatic cells of the cortex can be ZW or ZZ, with no intrinsic sex bias; the cells are naïve with respect to their chromosome sex identity.

Estrogen, normally provided by the female medulla, induces and maintains the formation of the cortex via activation of ERα within a sexually naïve epithelium. The finding that this activation can also occur above a male medulla, shows that estrogen is sufficient for this step and suggests that the initial formation of the cortex relies on positive inductive signals more than the lack of antagonising signals.

Later progression of the cortical germ cells into meiosis requires a medulla of the correct phenotypic sex, suggesting that meiotic entry is a checkpoint of a well-coordinated ovarian cortex/medulla differentiation pathway.

Our results provide new insights into the process of chicken ovarian differentiation and provide a model that could be extended to other systems, including many mammals. Moreover, our data suggest that the chick intersex models may be a potential valuable system for investigating the aetiology of germ cell tumours.

## Material and Methods

### Animals

Most experiments were performed using Dekalb white chicken eggs obtained from Henry Stewart, Louth, UK, except for the generation of chimeras, done using GFP transgenic chickens (Roslin greens) and ISA brown chickens held at the National Avian Research Facility (NARF) at Roslin Institute, Edinburgh, UK.

### Plasmids and *in ovo* electroporation

The Tol2 conditional integration plasmid system was a kind gift of dr. Yoshiko Takahashi (Kyoto, Japan) and includes: the transposase expression plasmid pCAGGS-T2TP, the conditional expression plasmid PT2K-B1-TRE-EGFP (carrying an EGFP reporter), the Tet-ON activator plasmid pT2K-CAG-rTA-M2 which responds to doxycycline (Watanabe et al., 2007). A truncated isoform of *ERα* lacking the last 196bp of the open reading frame (65aminoacids) was generated by PCR using primers F-5’ACTAGTTCTAGAGCCATTAGCAATGACCATGAC and R-5’GTCGACTCTAGATTAACACTTCATATTGTACAGGTG, cloned in pCRII-TOPO (Invitrogen) and then moved to PT2K-B1-TRE-EGFP (using BamHI and EcoRV sites) to make *cERα524*. This isoform corresponds to a human ERα isoform shown to have a powerful dominant negative activity (Ince et al., 1993).

Three *ERα* shRNA molecules were cloned downstream of a mouse U6 promoter into pCRIITOPO using a PCR based approach aiming to amplify the U6 promoter along with the shRNA, similar to the approach described in (Harper et al., 2005). Common forward primer: U6-Forward, 5’TCTAGATCGACGCCGCCATCTCTAG; U6-reverse-shRNA specific primers:

(ER1)CTCGAGAAAAAAGGAACACCCAGGAAAGCTTTCCCATCTGTGGCTTTACAGAAAGCTTTCCTGGGTGTTCCAAACAAGGCTTTTCTCCAAGGG;

(ER2)CTCGAGAAAAAAGCCACTAACCAGTGTACTATCCCATCTGTGGCTTTACAGATAGTACACTGGTTAGTGGCAAACAAGGCTTTTCTCCAAGGG,

(ER3)CTCGAGAAAAAAGGTACCCTACTACCTTGAAACCCATCTGTGGCTTTA

CAGTTTCAAGGTAGTAGGGTACCAAACAAGGCTTTTCTCClAAGGG.

The sequences include restriction enzymes for cloning. Dominant negative activity of cERα524 was confirmed by luciferase assays in vitro, following transfection in HEK293 cells together with full length *cERα* (cloned into pSG1) and a 3XERE-TATA-Luc luciferase reporter (Lemmen et al., 2004), kind gift of dr. Paul T. van der Saag, Ubrecht Institute, Utrecht, Netherlands. The three shER RNA molecules were also tested by luciferase assay using the same in vitro system. U6-shER ER1, displaying the best silencing activity were later cloned into pSLAX13, previously modified to carry IRES-EGFP. U6-shER-IRES-EGFP (ER1) were then moved to a RCAS(A) retroviral vector to generate RCAS-U6-shER-IRES-EGFP (RCAS-shER1).

*In ovo* electroporation of plasmid DNA into the left gonadal epithelium was achieved by DNA injection into the left coelomic cavity of HH15-17 embryos using a glass capillary needle and an Inject+matic pico pump, followed by electroporation using a NEPA21 electroporator (Sonidel) (five 50ms pulses at 26V, transferring pulse only). Detailed procedure as previously described (Guioli et al., 2007).

### *In ovo* drug treatment

For manipulation of estrogen levels after sex determination (late treatment), eggs were incubated at 37.7°C pointed end down. At D7-7.5 (HH31) or D9 (HH35) a first injection in the air chamber was performed through a hole made at the rounded end. To downregulate estrogen, the P450aromatase inhibitor Fadrozole (Sigma F3806) was injected at 0.5mg/egg in 50ul of PBS and then re-injected at 0.3mg/egg every other day. To increase estrogen, ß-Estradiol (Sigma E2758) was injected at 120µg/egg in 25µl of 95% ETOH; embryos collected at D14 (HH40) or D17 (HH43) were injected once more at D13 (HH39). For manipulation of estrogen levels before sex determination (early treatment) eggs were injected once at D4 (HH23) with 0.5mg or 1mg/egg of Fadrozole, or 120µg/egg of ß-estradiol.

### Generation of gonadal chimera

D2 (HH10/12) (13–15 somites) GFP transgenic embryos (Roslin greens) and ISA brown embryos were used as donor and host, respectively. Details of the procedure are described in (Zhao et al., 2010). The manipulated host embryos were then incubated until D18 (HH44).

### Antibodies, Immunohistochemistry and in situ hybridisation

The following antibodies were used: rat anti-VASA (1:1000) (Aramaki et al., 2009), rat anti ERα (1:200, Fitzgerald), rabbit anti-DMRT1 (1:1500) (Guioli and Lovell-Badge, 2007), goat anti-SOX9 (1:500, R&D), goat anti-FOXL2 (1:400, Abcam), mouse anti-P450aromatase (1:200, Serotec), mouse anti-γH2AX (1:200, Upstate), rabbit anti-SYCP3 (1:500, Novus), mouse anti-Fibronectin (1:100, Developmental Studies Hybridoma Bank), goat anti-LHX9 (1:200, Santacruz). The *STRA8* probe template for RNA *in situ* was generated by PCR cloning into PCRIITOPO (Invitrogen), using primers F-5’TACCCAGACACCTCATCCCC and R5’ TCAAAGGTCTCCGTGCACCG. The *PITX2* probe was previously described in (Logan et al., 1998). Urogenital ridges were fixed in 4% paraformaldehyde at 4°C overnight, rinsed in PBS at Room Temperature (RT), then transferred to 30% sucrose overnight and finally embedded in OCT and stored at −80°C. Cryosections for immunofluorescence were rinsed 3X five minutes in PBS and transferred for 1 hour to a blocking solution (PBS/0.1% triton, 2% donkey serum) before adding the primary antibody ON at 4°C (or 37°C for ERα). After 3X ten minute washes in PBS/0.1%tween, the sections were incubated with secondary antibodies (Invitrogen Alexa fluor-conjugated donkey antibodies 1:400 in PBS/0.1% tween). The results from the *in situ* hybridization on cryosections were imaged using a Leica DM-RA2 upright microscope equipped with a Retiga 200R camera and Q-capture Pro7 software (Q imaging). Fluorescence images were collected on a Olympus VS120 slide scanner equipped with a XM10 monochrome camera and a VS-ASW-L100 software (Olympus), or on a Leica upright SPE confocal system.

## Acknowledgments

We wish to thank dr. Yoshiko Takahashi for the Tol2 plasmid system; dr. Masa-aki Hattori for the VASA antibody; the Developmental Studies Hybridoma Bank (DSHB) at the University of Iowa, US for providing the Fibronectin antibody; dr. Paul T. van der Saag, for the luciferase reporter plasmid; dr. Thushyanthan Guruparan for his help with some electroporation experiments during his summer student internship; dr. Christophe Galichet for critical reading of the manuscript; Marie Caulfield and Jack Waterford in the biological research facility at the Francis Crick Institute for assistance; the light microscopy facility at the MRC National Institute for Medical Research and at the Francis Crick institute for technical support; the National Avian Research facility at the Roslin Institute for assistance.

## Author contributions

Conceptualisation: SG, RLB, MC; Methodology: SG, DZ; Validation: SG, DZ; Formal analysis: SG, DZ; Investigation: SG, DZ, SN; Visualisation: SG; Writing-original draft: SG; Writing-review and editing: SG, RLB, MC; Project administration: RLB, MC; Funding acquisition: RLB, MC.

## References

Akazome, Y. and Mori, T. (1999). Evidence of sex reversal in the gonads of chicken embryos after oestrogen treatment as detected by expression of lutropin receptor. J Reprod Fertil 115, 9–14.

Andrews, J. E., Smith, C. A. and Sinclair, A. H. (1997). Sites of estrogen receptor and aromatase expression in the chicken embryo. Gen Comp Endocrinol 108, 182–190.

Aramaki, S., Kubota, K., Soh, T., Yamauchi, N. and Hattori, M. A. (2009). Chicken dead end homologue protein is a nucleoprotein of germ cells including primordial germ cells. J Reprod Dev 55, 214–218.

Boulanger, L., Pannetier, M., Gall, L., Allais-Bonnet, A., Elzaiat, M., Le Bourhis, D., Daniel, N., Richard, C., Cotinot, C., Ghyselinck, N. B., et al. (2014). FOXL2 is a female sex-determining gene in the goat. Curr Biol 24, 404–408.

Bowles, J., Feng, C. W., Miles, K., Ineson, J., Spiller, C. and Koopman, P. (2016). ALDH1A1 provides a source of meiosis-inducing retinoic acid in mouse fetal ovaries. Nat Commun 7, 10845.

Bruggeman, V., Van As, P. and Decuypere, E. (2002). Developmental endocrinology of the reproductive axis in the chicken embryo. Comp Biochem Physiol A Mol Integr Physiol 131, 839–846.

Byskov, A. G. (1986). Differentiation of mammalian embryonic gonad. Physiol Rev 66, 71–117.

Carlon, N. and Stahl, A. (1985). Origin of the somatic components in chick embryonic gonads. Arch Anat Microsc Morphol Exp 74, 52–59.

Cools, M., Stoop, H., Kersemaekers, A. M., Drop, S. L., Wolffenbuttel, K. P., Bourguignon, J. P., Slowikowska-Hilczer, J., Kula, K., Faradz, S. M., Oosterhuis, J. W., et al. (2006). Gonadoblastoma arising in undifferentiated gonadal tissue within dysgenetic gonads. J Clin Endocrinol Metab 91, 2404–2413.

Coveney, D., Shaw, G. and Renfree, M. B. (2001). Estrogen-induced gonadal sex reversal in the tammar wallaby. Biol Reprod 65, 613–621.

de Melo Bernardo, A., Heeren, A. M., van Iperen, L., Fernandes, M. G., He, N., Anjie, S., Noce, T., Ramos, E. S. and de Sousa Lopes, S. M. (2015). Meiotic wave adds extra asymmetry to the development of female chicken gonads. Mol Reprod Dev 82, 774–786.

DeFalco, T. and Capel, B. (2009). Gonad morphogenesis in vertebrates: divergent means to a convergent end. Annu Rev Cell Dev Biol 25, 457–482.

Gasc, J. M. (1980). Estrogen target cells in gonads of the chicken embryo during sexual differentiation. J Embryol Exp Morphol 55, 331–342.

Ginsburg, M. and Eyal-Giladi, H. (1987). Primordial germ cells of the young chick blastoderm originate from the central zone of the area pellucida irrespective of the embryo-forming process. Development 101, 209–219.

Guioli, S. and Lovell-Badge, R. (2007). PITX2 controls asymmetric gonadal development in both sexes of the chick and can rescue the degeneration of the right ovary. Development 134, 4199–4208.

Guioli, S., Lovell-Badge, R. and Turner, J. M. (2012). Error-prone ZW pairing and no evidence for meiotic sex chromosome inactivation in the chicken germ line. PLoS Genet 8, e1002560.

Guioli, S., Nandi, S., Zhao, D., Burgess-Shannon, J., Lovell-Badge, R. and Clinton, M. (2014). Gonadal asymmetry and sex determination in birds. Sex Dev 8, 227–242.

Guioli, S., Sekido, R. and Lovell-Badge, R. (2007). The origin of the Mullerian duct in chick and mouse. Dev Biol 302, 389–398.

Hamburger, V. and Hamilton, H. (1951). A series of normal stages in the development of the chick embryo. J Morphol 88, 49–92.

Hamilton, K. J., Hewitt, S. C., Arao, Y. and Korach, K. S. (2017). Estrogen Hormone Biology. Curr Top Dev Biol 125, 109–146.

Harper, S. Q., Staber, P. D., He, X., Eliason, S. L., Martins, I. H., Mao, Q., Yang, L., Kotin, R. M., Paulson, H. L. and Davidson, B. L. (2005). RNA interference improves motor and neuropathological abnormalities in a Huntington’s disease mouse model. Proc Natl Acad Sci U S A 102, 5820–5825.

He, B., Lin, J., Li, J., Mi, Y., Zeng, W. and Zhang, C. (2012). Basic fibroblast growth factor suppresses meiosis and promotes mitosis of ovarian germ cells in embryonic chickens. Gen Comp Endocrinol 176, 173–181.

Hersmus, R., van Bever, Y., Wolffenbuttel, K. P., Biermann, K., Cools, M. and Looijenga, L. H. (2017). The biology of germ cell tumors in disorders of sex development. Clin Genet 91, 292–301.

Ince, B. A., Zhuang, Y., Wrenn, C. K., Shapiro, D. J. and Katzenellenbogen, B. S. (1993). Powerful dominant negative mutants of the human estrogen receptor. J Biol Chem 268, 14026–14032.

Ishimaru, Y., Komatsu, T., Kasahara, M., Katoh-Fukui, Y., Ogawa, H., Toyama, Y., Maekawa, M., Toshimori, K., Chandraratna, R. A., Morohashi, K., et al. (2008). Mechanism of asymmetric ovarian development in chick embryos. Development 135, 677–685.

Jorgensen, A., Nielsen, J. E., Almstrup, K., Toft, B. G., Petersen, B. L. and Rajpert-De Meyts, E. (2013). Dysregulation of the mitosis-meiosis switch in testicular carcinoma in situ. J Pathol 229, 588–598.

Karagenc, L., Cinnamon, Y., Ginsburg, M. and Petitte, J. N. (1996). Origin of primordial germ cells in the prestreak chick embryo. Dev Genet 19, 290–301.

Koubova, J., Hu, Y. C., Bhattacharyya, T., Soh, Y. Q., Gill, M. E., Goodheart, M. L., Hogarth, C. A., Griswold, M. D. and Page, D. C. (2014). Retinoic acid activates two pathways required for meiosis in mice. PLoS Genet 10, e1004541.

Lambeth, L. S., Ohnesorg, T., Cummins, D. M., Sinclair, A. H. and Smith, C. A. (2014). Development of retroviral vectors for tissue-restricted expression in chicken embryonic gonads. PLoS One 9, e101811.

Lemmen, J. G., Arends, R. J., van Boxtel, A. L., van der Saag, P. T. and van der Burg, B. (2004). Tissue- and time-dependent estrogen receptor activation in estrogen reporter mice. J Mol Endocrinol 32, 689–701.

Logan, M., Pagan-Westphal, S. M., Smith, D. M., Paganessi, L. and Tabin, C. J. (1998). The transcription factor Pitx2 mediates situs-specific morphogenesis in response to left-right asymmetric signals. Cell 94, 307–317.

Matson, C. K., Murphy, M. W., Griswold, M. D., Yoshida, S., Bardwell, V. J. and Zarkower, D. (2010). The mammalian doublesex homolog DMRT1 is a transcriptional gatekeeper that controls the mitosis versus meiosis decision in male germ cells. Dev Cell 19, 612–624.

McQueen, H. A. and Clinton, M. (2009). Avian sex chromosomes: dosage compensation matters. Chromosome Res 17, 687–697.

Nakamura, M., Kuwana, T., Miyayama, Y. and Fujimoto, T. (1988). Extragonadal distribution of primordial germ cells in the early chick embryo. Anat Rec 222, 90–94.

Omotehara, T., Smith, C. A., Mantani, Y., Kobayashi, Y., Tatsumi, A., Nagahara, D., Hashimoto, R., Hirano, T., Umemura, Y., Yokoyama, T., et al. (2014). Spatiotemporal expression patterns of doublesex and mab-3 related transcription factor 1 in the chicken developing gonads and Mullerian ducts. Poult Sci 93, 953–958.

Rodriguez-Leon, J., Rodriguez Esteban, C., Marti, M., Santiago-Josefat, B., Dubova, I., Rubiralta, X. and Izpisua Belmonte, J. C. (2008). Pitx2 regulates gonad morphogenesis. Proc Natl Acad Sci U S A 105, 11242–11247.

Schmidt, D., Ovitt, C. E., Anlag, K., Fehsenfeld, S., Gredsted, L., Treier, A. C. and Treier, M. (2004). The murine winged-helix transcription factor Foxl2 is required for granulosa cell differentiation and ovary maintenance. Development 131, 933–942.

Smith, C. A., Roeszler, K. N., Bowles, J., Koopman, P. and Sinclair, A. H. (2008). Onset of meiosis in the chicken embryo; evidence of a role for retinoic acid. BMC Dev Biol 8, 85.

Smith, C. A., Roeszler, K. N., Ohnesorg, T., Cummins, D. M., Farlie, P. G., Doran, T. J. and Sinclair, A. H. (2009). The avian Z-linked gene DMRT1 is required for male sex determination in the chicken. Nature 461, 267–271.

Tassinari, V., Campolo, F., Cesarini, V., Todaro, F., Dolci, S. and Rossi, P. (2015). Fgf9 inhibition of meiotic differentiation in spermatogonia is mediated by Erk-dependent activation of Nodal-Smad2/3 signaling and is antagonized by Kit Ligand. Cell Death Dis 6, e1688.

Uda, M., Ottolenghi, C., Crisponi, L., Garcia, J. E., Deiana, M., Kimber, W., Forabosco, A., Cao, A., Schlessinger, D. and Pilia, G. (2004). Foxl2 disruption causes mouse ovarian failure by pervasive blockage of follicle development. Hum Mol Genet 13, 1171–1181.

Uhlenhaut, N. H., Jakob, S., Anlag, K., Eisenberger, T., Sekido, R., Kress, J., Treier, A. C., Klugmann, C., Klasen, C., Holter, N. I., et al. (2009). Somatic sex reprogramming of adult ovaries to testes by FOXL2 ablation. Cell 139, 1130–1142.

Vaillant, S., Guemene, D., Dorizzi, M., Pieau, C., Richard-Mercier, N. and Brillard, J. P. (2003). Degree of sex reversal as related to plasma steroid levels in genetic female chickens (Gallus domesticus) treated with Fadrozole. Mol Reprod Dev 65, 420–428.

Watanabe, T., Saito, D., Tanabe, K., Suetsugu, R., Nakaya, Y., Nakagawa, S. and Takahashi, Y. (2007). Tet-on inducible system combined with in ovo electroporation dissects multiple roles of genes in somitogenesis of chicken embryos. Dev Biol 305, 625–636.

Yang, X., Zheng, J., Na, R., Li, J., Xu, G., Qu, L. and Yang, N. (2008). Degree of sex differentiation of genetic female chicken treated with different doses of an aromatase inhibitor. Sex Dev 2, 309–315.

Yu, M., Yu, P., Leghari, I. H., Ge, C., Mi, Y. and Zhang, C. (2013). RALDH2, the enzyme for retinoic acid synthesis, mediates meiosis initiation in germ cells of the female embryonic chickens. Amino Acids 44, 405–412.

Zarkower, D. (2013). DMRT genes in vertebrate gametogenesis. Curr Top Dev Biol 102, 327–356.

Zhao, D., McBride, D., Nandi, S., McQueen, H. A., McGrew, M. J., Hocking, P. M., Lewis, P. D., Sang, H. M. and Clinton, M. (2010). Somatic sex identity is cell autonomous in the chicken. Nature 464, 237–242.

