## Supplementary Figures for "In the chick embryo, estrogen can induce chromosomally male ZZ left gonad epithelial cells to form an ovarian cortex, which supports oogenesis"

**Supplementary material**

**Figure legends**

**Supplementary Fig. 1 In ZW embryos treated with Fadrozole at D7-7.5 (HH31) the growth of the left ovary is affected.** (A) D17 (HH43) ZW control gonadal pair; (B) ZW gonadal pair from embryos treated with Fadrozole from D7-7.5 (ZW-Fa); (C, D) Fluorescent images of cryostat sections from the left gonad in (A) and (B) respectively, stained for the female markers P450aromatase (red) and FOXL2 (green). ZW-Fa left gonads have impaired overall growth but retain female identity. White dotted lines highlight the cortex-medulla border.

**Supplementary Fig. 2 The left testis of ZZ embryos exposed to ß-estradiol at D9 (HH35) develops a cortical domain.** Sections from the left gonad of D17 (HH43) embryos, immunostained for the Sertoli cell marker SOX9 (red) and the germ cell marker P63 (green). (A) ZZ control, (B) ZW control, (C) ZZ treated with ß-estradiol (ZZ-E2) at D9 (HH35). ZZ left gonads remain sensitive to ß-estradiol during testis differentiation at least until embryonic D9. White dotted lines highlight the cortex-medulla border.

**Supplementary Fig. 3 ERa protein is produced in the epithelium of the left ZZ gonad during differentiation.** Sections from ZW and ZZ left and right wild type gonads at embryonic D7 (HH31), D8 (HH34) and D12 (HH38), immunostained for ERα (red). ER α is expressed in both ZW female and ZZ male left gonadal epithelium at sex determination and during sex specific differentiation. White dotted lines in left gonad panels highlight the cortex-medulla border.

**Supplementary Fig. 4 *In ovo* suppression of epithelial ERα by RNA interference (RNAi) disrupts cortex differentiation.** (A) Schematic of the construct: RCAS retroviral vector expressing *ERα* specific short hairpin RNA molecules in tandem with an EGFP reporter (RC-shER). (B) Whole mount image of a ZW left gonad electroporated with RC-shER1 at D2.5 (HH15-17) and screened at embryonic D9.5 (HH35-36). The EGFP reporter (green) marks the targeted cells. (C-F) Fluorescent images of sections from D9.5 (HH35-36) left gonads immunostained for P450 aromatase (P450) (red) or VASA (red); (C, E) ZW female control; (D-F) ZW RC-shER1 gonad shown in (B). Downregulation of epithelial ERα via RNAi compromises the cortex domain without affecting medulla sex identity. White dotted lines highlight the the cortex borders.

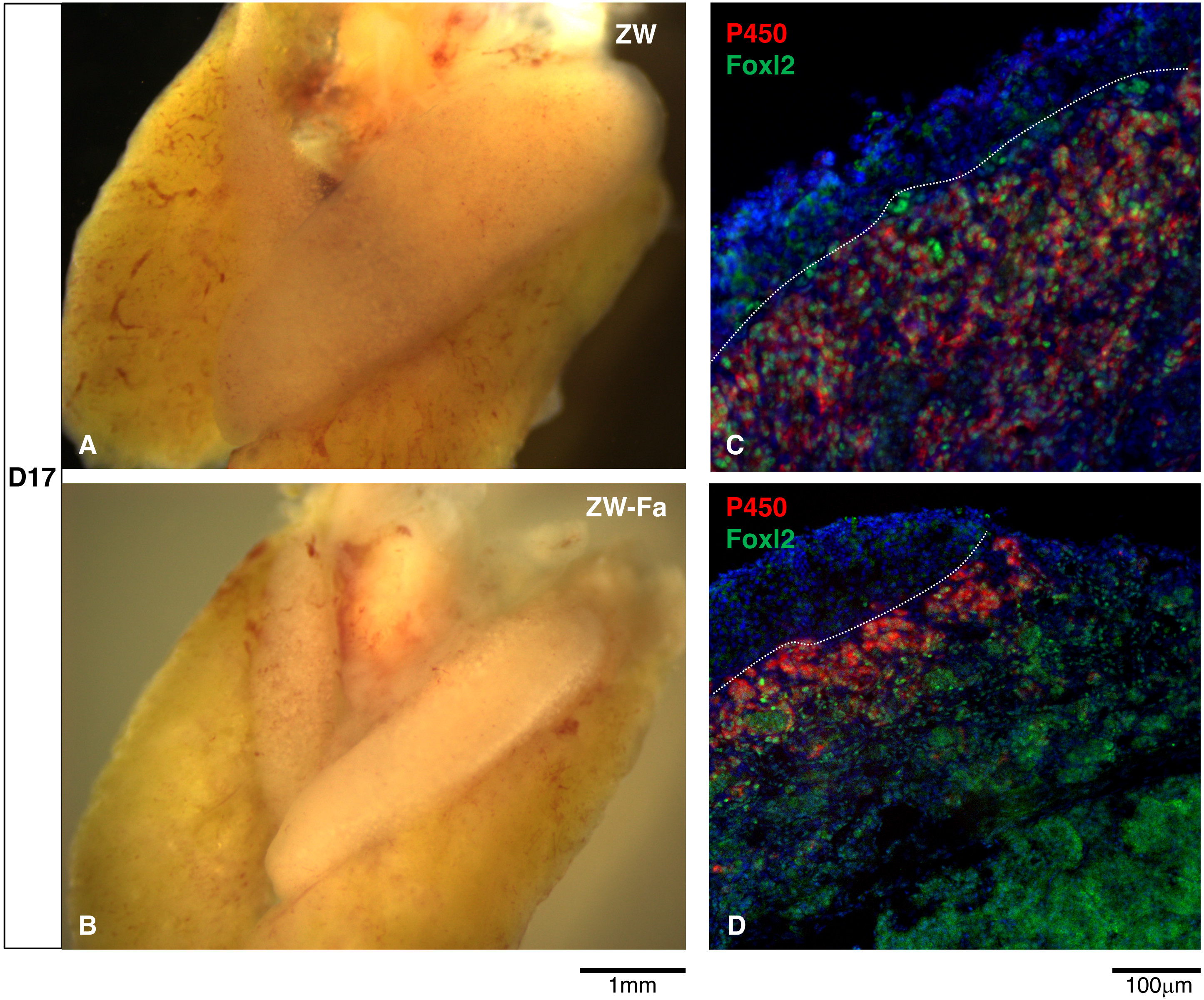


Suppl. Fig.1

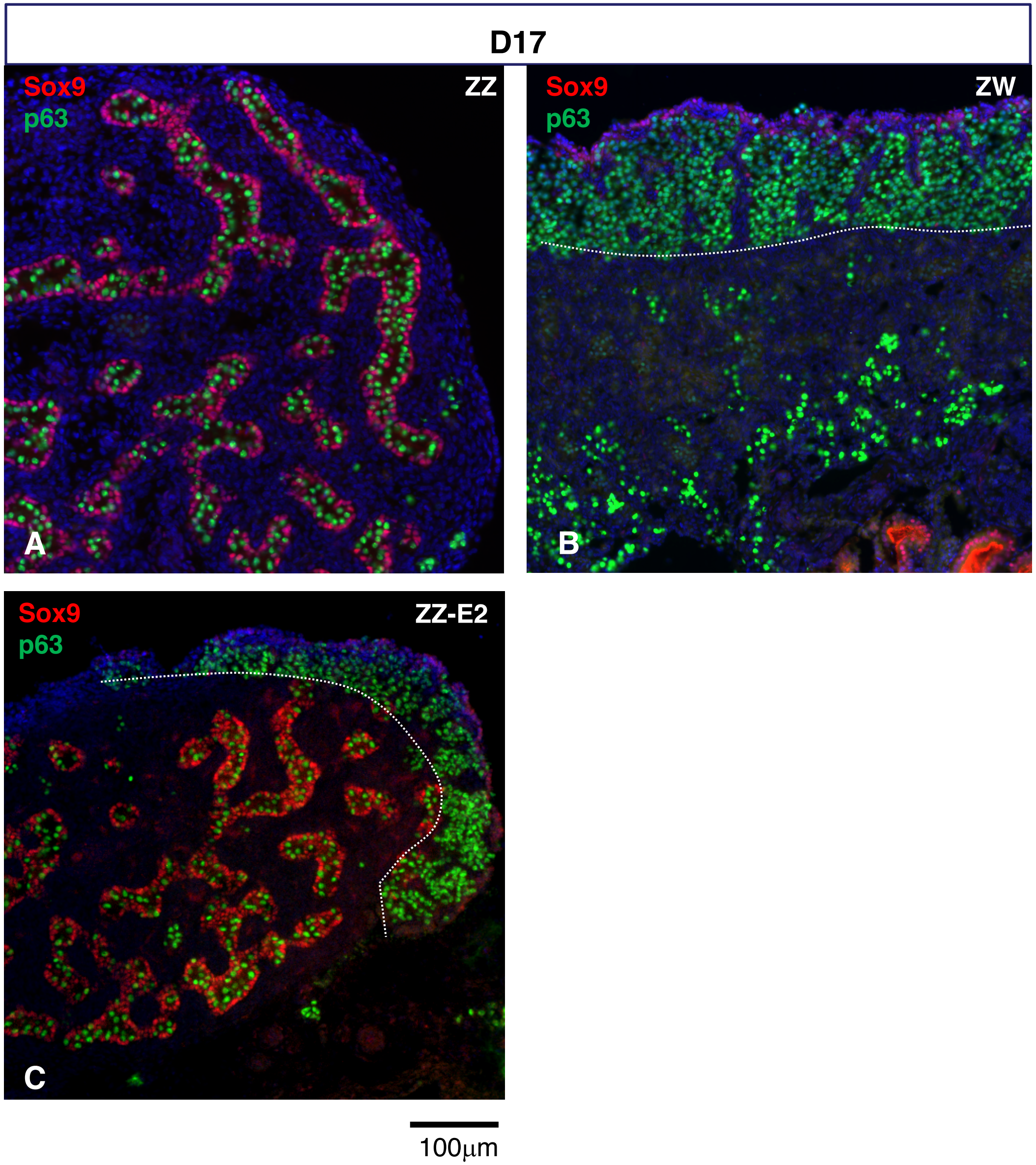


Suppl. Fig.2

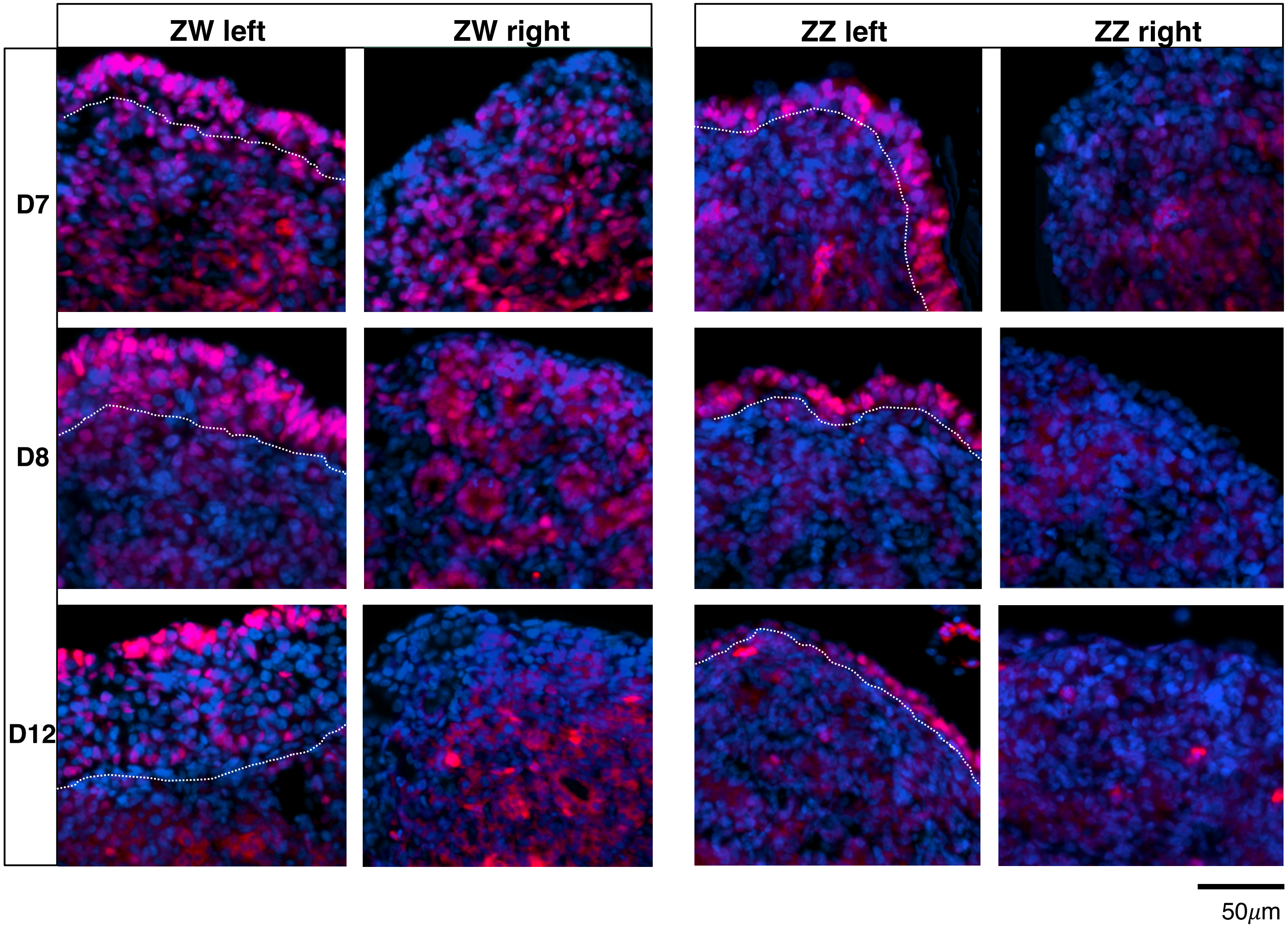


Suppl. Fig.3

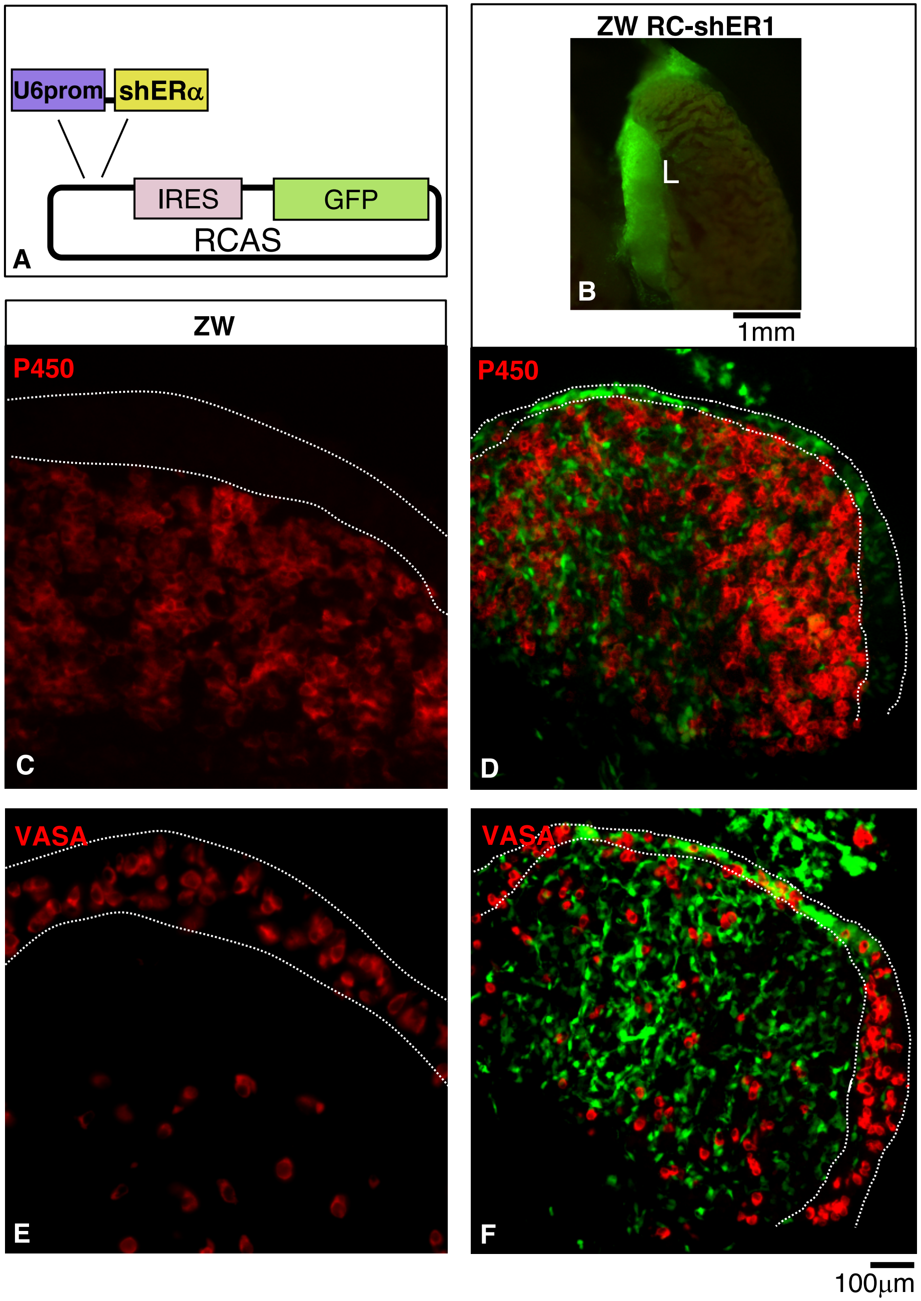
U6prom
shERα
IRES
GFP
RCAS
A
ZW RC-shER1
L
B
1mm
ZW
P450
C
P450
D
VASA
E
VASA
F
100µm

Suppl. Fig. 4
